# Phylogeographic separation and formation of sexually discrete lineages in a global population of *Yersinia pseudotuberculosis*

**DOI:** 10.1101/149468

**Authors:** Tristan Seecharran, Laura Kalin-Mänttäri, Katja A. Koskela, Simo Nikkari, Benjamin Dickins, Jukka Corander, Mikael Skurnik, Alan McNally

## Abstract

*Yersinia pseudotuberculosis* is a Gram negative intestinal pathogen of humans and has been responsible for several nation-wide gastro-intestinal outbreaks. Large-scale population genomic studies have been performed on the other human pathogenic *Yersinia, Y. pestis* and *Y. enterocolitica* allowing a high-resolution understanding of the ecology, evolution and dissemination of these pathogens. However, to date no large-scale global population genomic analysis of *Y. pseudotuberculosis* has been performed. Here we present analyses of the genomes of 134 strains of *Y. pseudotuberculosis* isolated from around the world, from multiple ecosystems since 1960’s. Our data display a phylogeographic split within the population, with an Asian ancestry and subsequent dispersal of successful clonal lineages into Europe and the rest of the world. These lineages can be differentiated by CRISPR cluster arrays, and we show that the lineages are limited with respect to inter-lineage genetic exchange. This restriction of genetic exchange maintains the discrete lineage structure in the population despite co-existence of lineages for thousands of years in multiple countries. Our data highlights how CRISPR can be informative of the evolutionary trajectory of bacterial lineages, and merits further study across bacteria.

## Introduction

The genus *Yersinia* belongs to the Gram-negative Enterobacteriaceae family of bacteria, and is a model genus for studying the evolution of bacterial pathogens ^1^. Three species of *Yersinia* are well-recognised human pathogens: the plague bacillus *Y. pestis*, and the enteropathogenic *Y. pseudotuberculosis*, and *Y. enterocolitica* ^1^. *Y. pseudotuberculosis*, which causes infection in a broad range of hosts, including domesticated and wild animals, has also been associated with foodborne infection in humans – known as yersiniosis. Transmission of the bacterium is usually through the faecal-oral route, and human infection can result from the ingestion of contaminated food products or water, or otherwise by direct contact with an infected animal or human ^2–5^. *Y. pseudotuberculosis* is also found widely in the environment, including soil ^6^, and in animals it causes tuberculosis-like disease ^6^. Human cases of *Y. pseudotuberculosis* infections are usually sporadic, however several large outbreaks have been reported in Finland and recently in New Zealand ^7,8^. Classical identification and typing of *Y. pseudotuberculosis* is based on the lipopolysaccharide O-antigen, resulting in a total of 21 known serotypes ^9^. However the efficacy of serotyping is very limited due to a large proportion of strains belonging to serotypes O:1a, O:1b, and O:3 ^8,10^.

The population structure of *Y. pseudotuberculosis* has been elucidated by MLST ^10^. This added further granularity to the serotype differentiation, grouping serotype O:3 strains into a distinct clone designated ST19 which are characterised by a truncation in the Yersiniabactin locus resulting in the loss of the genes encoding Iron transport across the bacterial membrane. Serotype O:1 strains formed a distinct clade of strains encompassing a large number of sequence type complexes, suggesting a highly diverse population of bacteria within the serotype O:1 group of *Y. pseudotuberculosis*. In addition to MLST genotyping, recent work has also analysed clustered regularly interspaced short palindromic repeat (CRISPR) loci of 335 *Y. pseudotuberculosis* isolates ^11^. The CRISPR-Cas system is an RNA-based immune system that regulates invasion of plasmids and viruses in bacteria and archaea ^12^. CRISPRs are constructed from a chain of 21 to 47 bp repeated sequences [called direct repeats (DR)] and in between DRs are unique spacer sequences. These spacers represent foreign DNA originating predominantly from bacteriophages and plasmids. In *Y. pseudotuberculosis* the CRISPR spacers are stored in 3 genomic loci named YP1, YP2 and YP3, and we identified 1902 distinct spacer sequences. One central finding was that *Y. pseudotuberculosis* and *Y. pestis* strains shared very few spacers, and that *Y. pestis* carries a relatively low number of spacers compared with *Y. pseudotuberculosis*. This would indicate that differentiation in CRISPR profile occurs after the phylogenetic split due to differences in ecological habitat encountered by the bacteria.

To date the most comprehensive genome-scale analysis of *Y. pseudotuberculosis* centred around a country-wide outbreak in New Zealand ^8^. Incorporation of publically available genomes into this data set also suggested a highly diverse species and that the New Zealand strains represented a geographically isolated clade of *Y. pseudotuberculosis*. The paucity of a specifically designed geographically and temporally distributed dataset of *Y. pseudotuberculosis* genomes means that our understanding of the population structure and evolutionary events occurring within this species is poorly informed. Global phylogenomic studies of *Y. pestis* have identified evolution of a clone from *Y. pseudotuberculosis* as a result of gene loss and then global dissemination ^1,13^. In contrast such studies in *Y. enterocolitica* have pointed to evolutionary path from a non-pathogenic ancestor via gene gain and loss, resulting in apparently ecologically separated clades within the species ^14,15^. By analysing a set of geographically and temporally distributed genomes we show that evolution within *Y. pseudotuberculosis* differs from that seen in the other two human pathogenic *Yersinia* species. We report a previously undetected geographic split between Asian and European strains and the presence of discrete phylogenetic clusters within the species which correlate with specific patterns of CRISPR spacer cassettes. This CRISPR signature correlates with patterns of accessory gene sharing within the species as well as core genome recombination.

## Results

### Phylogeographic structure of *Y. pseudotuberculosis* signals an Asian ancestry

A maximum likelihood phylogeny was reconstructed from a core genome alignment of 134 *Y. pseudotuberculosis* genome sequences (Figure 1). The phylogeny has a clear two-clade structure with a seemingly ancestral clade containing high diversity and long branch lengths, and a second clade containing much lower levels of diversity. Annotation of the tree with geographical source of isolation identifies a very clear geographic split in the phylogeny, with the ancestral highly diverse clade containing primarily Asian isolates and the second low diversity clade containing primarily European isolates. Between these two clades is a small transitional cluster of isolates originating from South Africa, and North and South America. Such a phylogenetic structure is consistent with an Asian ancestry for *Y. pseudotuberculosis*, with two separate migrations into Africa and the Americas, and into Europe. A European migration is consistent with a bottleneck event leading to successful establishment of a small number of clones. Annotation of the phylogeny with serotype and host species (Fig S1) identified that the European clade is further split into serotype 1a and serotype 1b clusters, and that there is no pattern of phylogenetic grouping associated with host.

**Figure 1.**
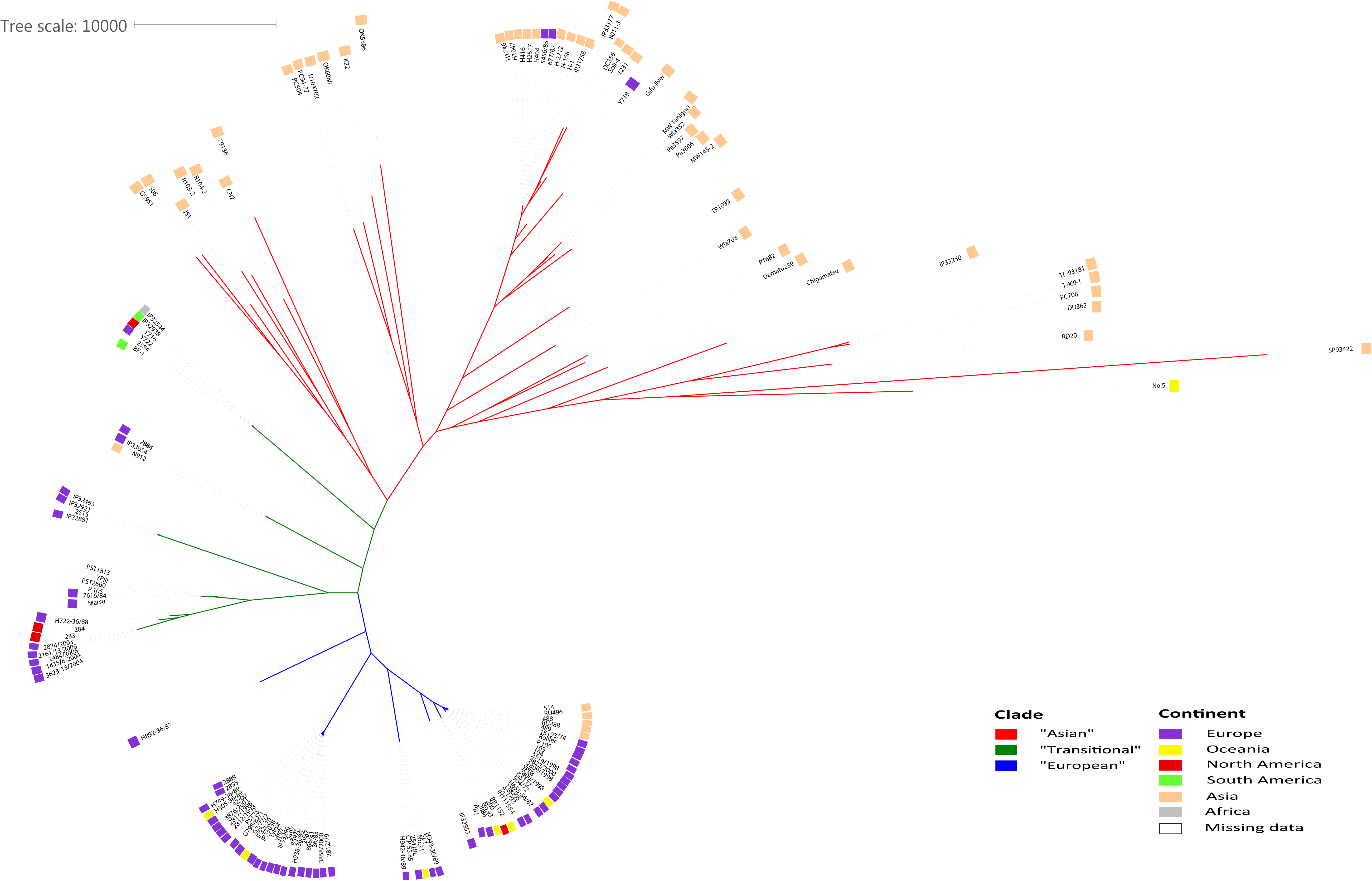
Maximum likelihood Phylogenetic tree of 134 *Y. pseudotuberculosis* isolates. The phylogeny is derived from a core genome alignment constructed using Parsnp and the phylogenetic tree was visualised using iTOL. Geographical source of isolation is superimposed on the tree as coloured bars. The “Asian” and “European” clades are defined by tree branch colouring.

### Phylogenetic dating suggests recent geographical divergence

Of the 134 genomes sequenced isolation dates are available for 73 isolates which represent the full diversity of the phylogeny. To date the evolutionary split of the “European” clade of strains from the “Asian” clade we used BEAST, analysing only the 73 strains for which isolation dates were available (Fig S2). The analysis suggests a tMRCA for the data set of 33,591 years before present (95% CI 49,460 – 20,476), with the divergence of the European and Asian clades occurring approximately 12,500 years ago. This period marks the transition between the Neolithic and Mesolithic eras at the end of the last ice age, and the beginning of livestock domestication and wheat and barley farming. A Bayesian skyline plot analysis of the dated phylogeny also supports the possibility of a strong bottleneck occurring in the population within the European clade, occurring in the very recent past (Fig S2).

### Phylogenetic clusters within *Y. pseudotuberculosis* associate with discrete CRISPR cassette patterns

We sought to determine any obvious genotypic traits associated with the phylogeographic split in our population. Bayesian clustering of the presence/absence of all 2969 known *Y. pseudotuberculosis* CRISPR spacers present in the data set identified a total of 33 distinct clusters of CRISPR spacer cassettes (Table S1). Annotation of these CRISPR clusters on the phylogenetic tree shows that the clusters form phylogenetically distinct groups of *Y. pseudotuberculosis* (Figure 2). The most recent of these clusters has a tMRCA to the rest of the population of 5,222 years before present, suggesting that this clustering is not a recent phenomenon nor is it due to any temporal artefacts of sampling. Indeed comparison of the CRISPR cluster pattern to year and geographical source of isolation (Table S1) suggests that this clustering is not a result of strains isolated in the same short time span or localised source. To confirm this we mapped the geographical source of isolation of all of the CRISPR clusters (Figure 3). This shows that CRISPR clusters are widely distributed across the world with some correlation to the higher phylogeographic split observed earlier. It also shows the highest diversity in CRISPR clusters occurs in Asia consistent with an Asian ancestry of *Y. pseudotuberculosis*.

**Figure 2.**
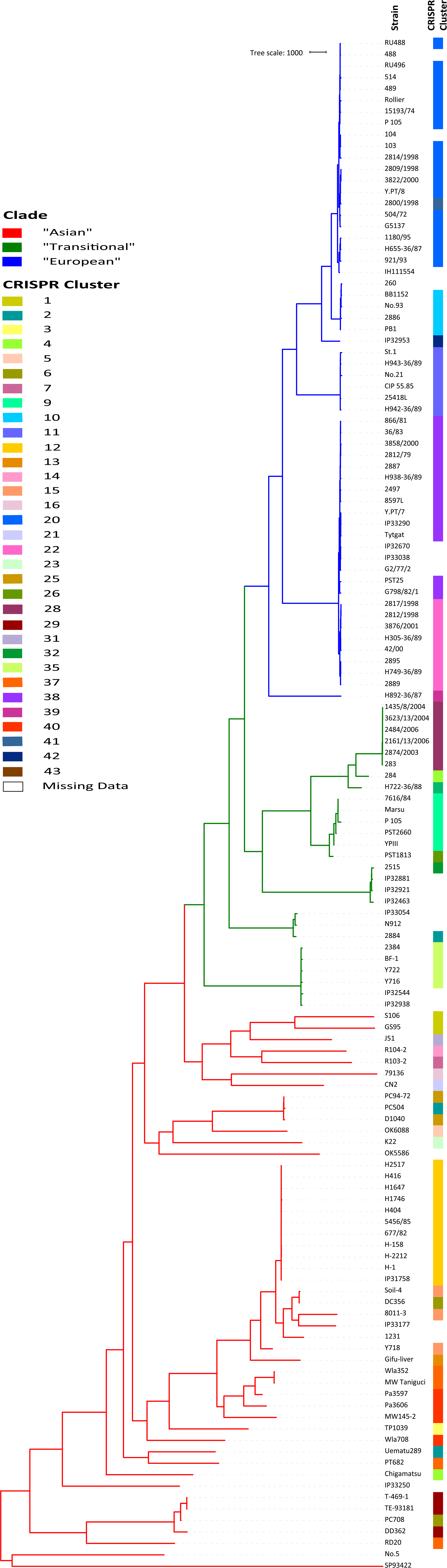
Phylogeny of 134 *Y. pseudotuberculosis* isolates annotated with CRISPR clusters as determined by Bayesian clustering of concatenated sequence of CRISPR spacer arrays. The “Asian” and “European” clades are defined by tree branch colouring.

**Figure 3.**
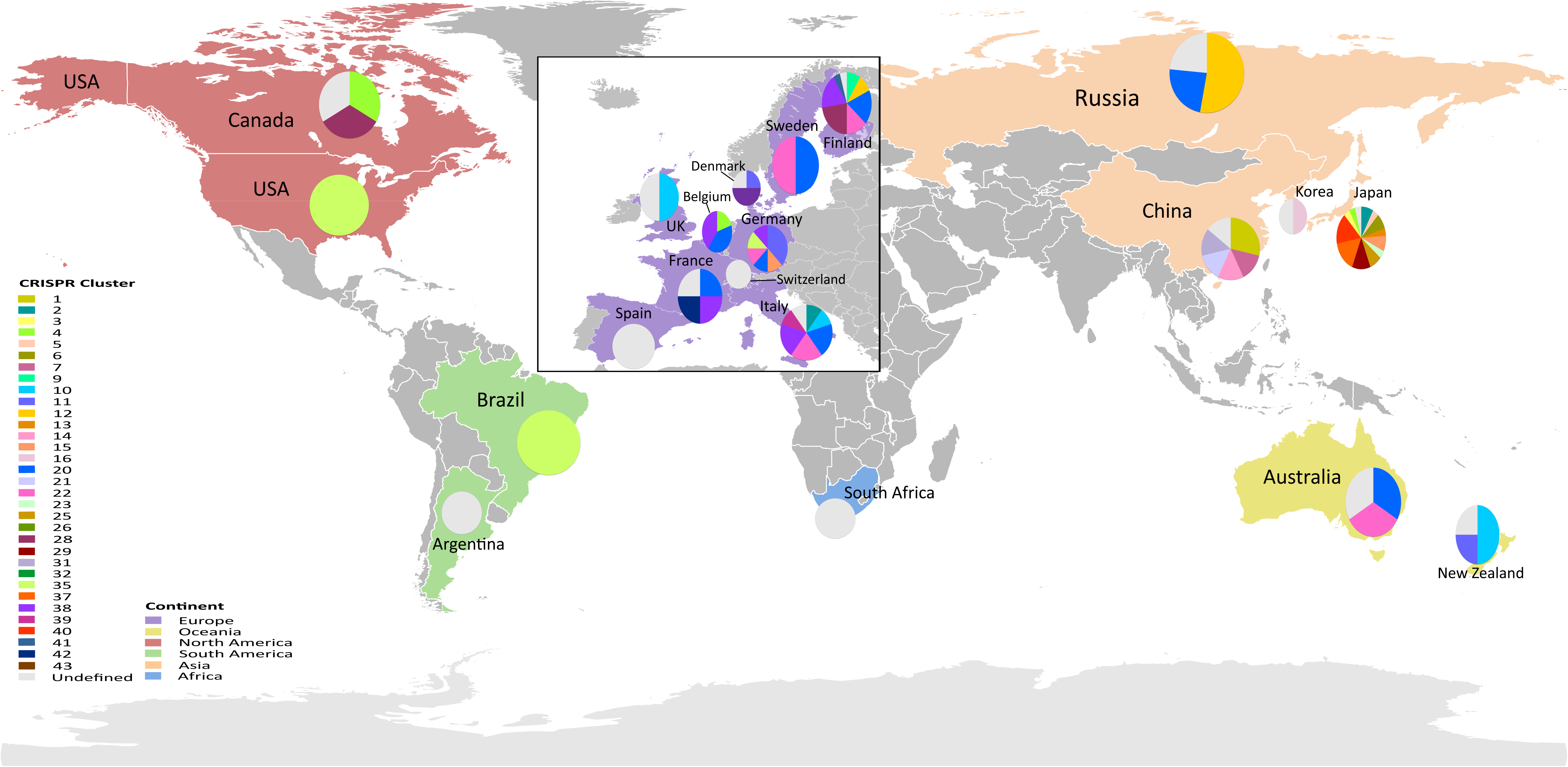
Global map showing the geographical sources of isolation of strains belonging to each of the 33 identified CRISPR clusters. The map shows clear co-existence of multiple CRISPR cluster-type strains in a number of countries

### Phylogenetic clusters are associated with patterns of genetic recombination in *Y. pseudotuberculosis.*

Given the role of CRISPR in generating lasting immunity to foreign DNA, we sought to investigate if the phylogenetic clusters within *Y. pseudotuberculosis* were associated with any signature of gene sharing. We created a pangenome matrix for all 134 genomes using LS-BSR, and then extracted the accessory genome. This accessory genome matrix was then used to annotate the core phylogenetic tree alongside CRISPR clusters (Figure 4). There are clear patterns of accessory genome profile concordant with the pattern of CRISPR clusters within the tree. Attempts to identify genes unique to any given phylogenetic cluster were largely unsuccessful. Only one unique coding sequence was detected in the “European” clade of strains and no unique CDSs in the “Asian” clade of strains. Of CRISPR clusters that contained more than one representative strain, CRISPR 22 contained nine unique CDSs. Other clusters include CRISPR 28 with two unique CDSs and CRISPRs 1, 9, and 11 each with one unique CDS, when compared to the rest of the population. This analysis suggests that each cluster contains a unique combination of accessory genes rather than unique genes per se. To confirm this, we calculated the average accessory genome dissimilarity for all CRISPR clusters containing more than one strain. This showed that in 12 of 18 clusters, strains have significantly more similar gene profiles to strains in the same cluster than to strains in other clusters (p<0.05 based on a 10000 random permutations test). This suggests that gene sharing between strains in the *Y. pseudotuberculosis* population is largely restricted to within individual phylogenetic clades. Analysis of core genome recombination identified a distribution of core genome recombination events which is highly concordant with the CRISPR clusters (Figure 5). Despite very high levels of recombination across the data set, the recombination occurring is not eroding the phylogenetic or CRISPR cluster signal, suggesting that inter-cluster horizontal transfer of genetic material is largely inhibited or occurs at very low frequency compared to intra-cluster recombination events.

**Figure 4.**
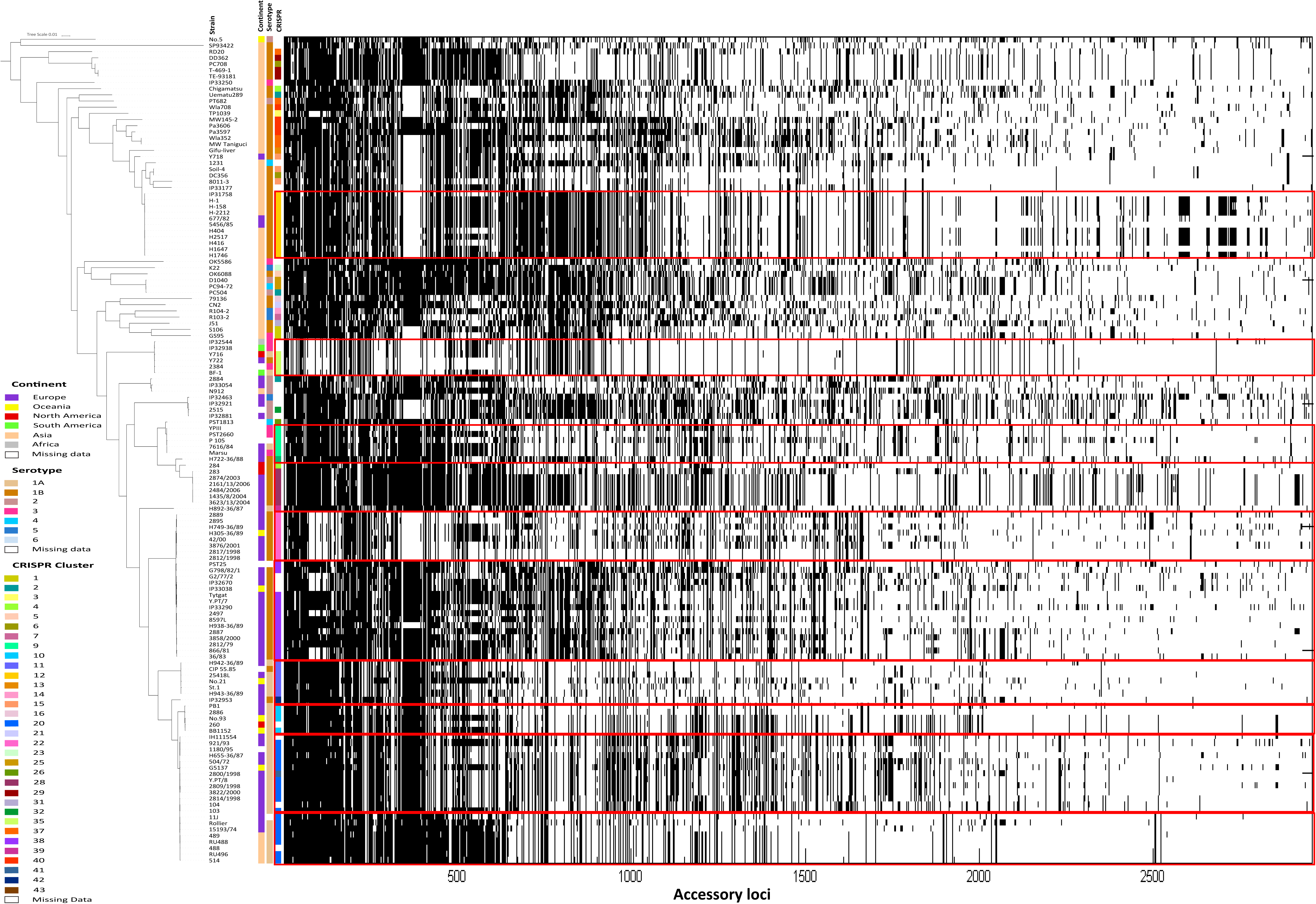
Distribution of accessory gene profiles for 134 *Y. pseudotuberculosis* isolates. The genes (columns) have been sorted by their presence/absence pattern (black, present; white, absent) across strains (rows), which have been sorted according to the phylogenetic tree. CRISPR clusters, continent and serotype are shown as colour bars.

**Figure 5.**
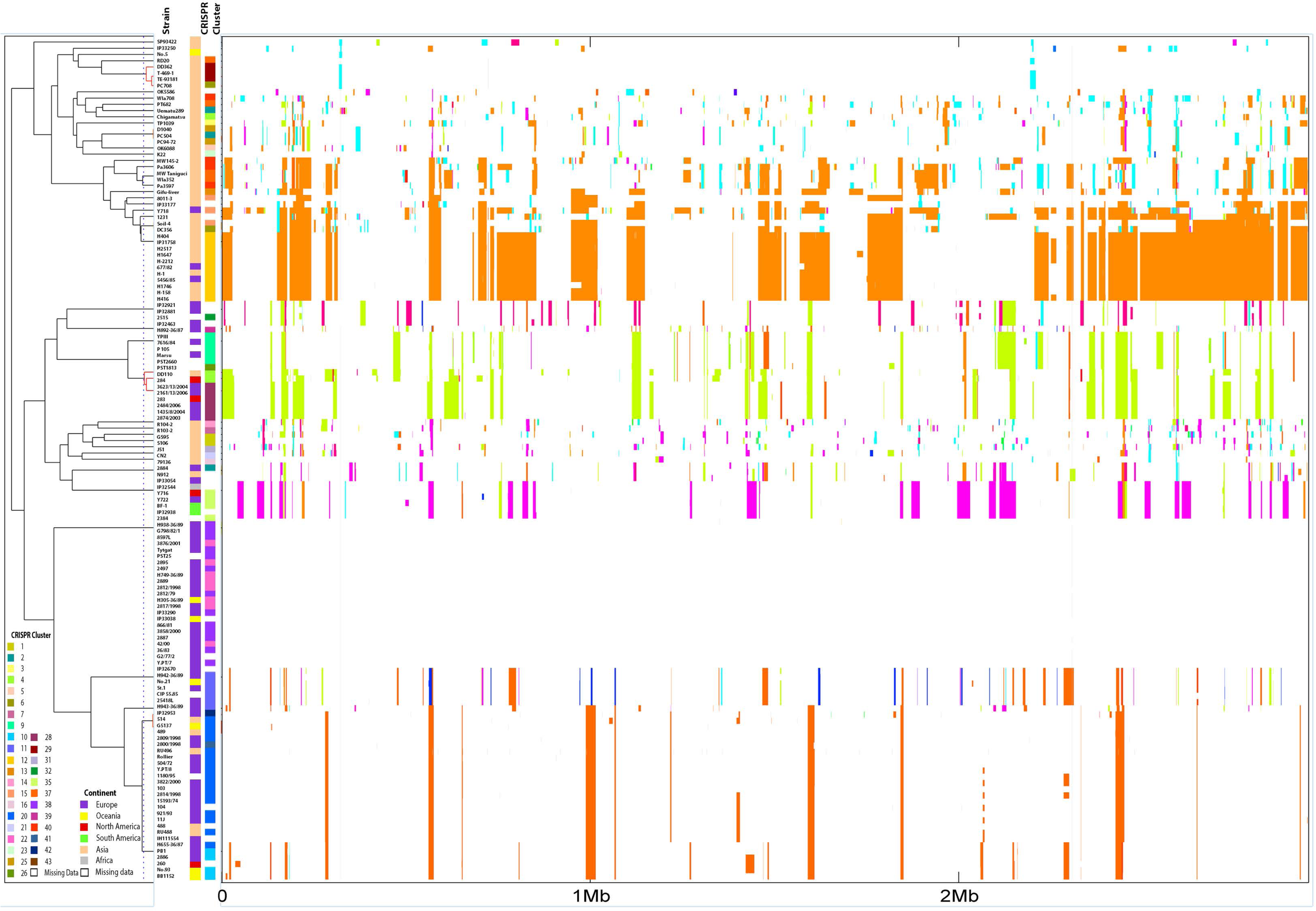
BratNextGen analysis of core genome recombination events for 134 *Y. pseudotuberculosis* isolates. Horizontal coloured bars show the indicated recombination events for each strain (y-axis); at the relative base pair positions (x-axis). CRISPR clusters and continent of origin are shown as colour bars. Where segments are the same colour indicates recombination events that are shared between those strains.

## Discussion

The genus *Yersinia* has acted as a model for developing our understanding of microbial pathogenesis, molecular microbiology, and microbial evolution ^1^. *Yersinia* was the first bacterial genus to have all representative species sequenced allowing fine scale analysis of how pathogenesis evolved in the three human pathogenic members of the genus ^15^. This analysis showed a striking degree of parallelism in how human-pathogenesis evolved in the pathogenic *Yersinia* ^15^. However finer scale evolutionary genomics studies of *Y. pestis* and *Y. enterocolitica* have shown very divergent mechanisms of intra-species evolution. *Y. pestis* is a recently evolved clone of *Y. pseudotuberculosis* which is globally disseminated and host-restricted with very low levels of diversity allowing fine scale transmission events to be successfully reconstructed ^13^. In complete contrast to this, pathogenic *Y. enterocolitica* have evolved from a non-pathogenic ancestor and have split into ecologically distinct clades which move rapidly across host species ^14^.

By sequencing a globally and temporally distributed set of 134 *Y. pseudotuberculosis* genomes isolated from a wide range of hosts and environments, we show that evolution in this species is driven by completely different mechanisms again. Our data shows that *Y. pseudotuberculosis* is the only pathogenic *Yersinia* species which shows a clear phylogeographic split in its population. This was once postulated to be the case for *Y. enterocolitica* ^16^ with Old World and New World strains, however comprehensive population genomics have shown this is not the case ^14,15^. The discovery of Asian ancestry in *Y. pseudotuberculosis* is in line with the postulated ancestry of *Y. pestis* ^13,17^, though our data appears to show the most outstanding genetic variation occurring in Japan, not China. Whilst this does not appear to be an artefact of sampling in this study it cannot be discounted that a more thorough and dense genomic sampling would provide a different result. However of interest is the fact that a sub-clade of *Y. pseudotuberculosis* exists which causes far-east scarlet-like fever and is associated with Japan and tropical South East Asia ^18^, suggesting larger variation in this region and a potential focus of ancestry for the species. Although accurate dating from a relatively small timed sample set is difficult, our tMRCA for the entire *Y. pseudotuberculosis* data set is in the same range (10K – 40K ya) as that postulated for the emergence of *Y. pestis* ^17^, and it is tempting to speculate that this emergence coincided with a larger population and dispersal event across *Y. pseudotuberculosis*.

Previous work by our group analysed patterns of accessory gene sharing to conclude that *Y. enterocolitica* was formed of ecologically distinct phylogroups ^14^. This hypothesis was formed on the basis that the limited inter-clade sharing of genes could not be due to stearic hindrance by O-antigen nor be genetic exclusion as no such mechanisms existed. Our data on *Y. pseudotuberculosis* also identifies clearly distinct phylogroups within the phylogeographic clades. These phylogroups have unique combinations of accessory genes with little variation in the accessory genome, and a very similar pattern of core genome homologous recombination. Similar to *Y. enterocolitica* it is highly improbable that this might be driven by some factor which precludes physical contact given the limited serotypes present in *Y. pseudotuberculosis* ^4^. Rather our analysis is strongly indicative of genetic restriction between phylogroups, and that this can be correlated with patterns of acquired CRISPR cassettes. The primary evidence for this genetic restriction is in the fact that different phylogroups co-exist in the same geographical locations. In the absence of any active barrier to recombination one would expect the signal that identifies each CRISPR cluster to be eroded relatively quickly in time ^19^ resulting in a lack of clear phylogroup signatures ^20^. This would be especially so given the large levels of recombination detected in the core genome of *Y. pseudotuberculosis*. As the phylogenetic clusters have co-existed in locations for around 5,000 years or more and continue to display a clear signal of within cluster similarity, our data suggest very limited genetic exchange between clusters and maintenance of the distinct phylogroups.

CRISPR has been shown to play a role in dictating the accessory genomes of *Pseudomonas aeruginosa* ^21^, and CRISPR analysis correlated with phylogenetic structure in a study of *Shigella* genomes ^22^. Previous data has shown that CRISPR evolution in bacteria, and particularly in the Enterobacteriaceae is driven by vertical and not horizontal evolution ^23^. Together our data create a hypothesis for *Y. pseudotuberculosis* evolution whereby large population perturbations give rise to geographically isolated clones. During the early formation of these clones exposure to geographically localised exogenous DNA creates a CRISPR array of immunity which correlates with the repertoire of genetic material that can be transferred and acquired from the gene pool. As each of these clones then globally disseminates they find themselves in co-existence with other clones of *Y. pseudotuberculosis*, but gene transfer between clones is restricted. This restriction is such that transfer of genetic material cannot occur at levels required to erode the clonal phylogenetic signature in the population, and consequently distinct phylogroups of *Y. pseudotuberculosis* persist in the population.

## Methods

### Bacterial isolates and genome sequences

A total of 134 *Y. pseudotuberculosis* genomes were analysed in this study, of which 108 were newly sequenced (Table S1). These isolates were collected over a 46-year time frame from a wide host range covering 19 different countries and 6 continents, and represent the full spectrum of serotypes possible. Additionally the strains were isolated from a wide range of hosts including human clinical, livestock, wild animals, companion animals, and environmental sources. Library preparation and sequencing of these isolates were performed using the Illumina Nextera kit and Genome Analyzer IIx instrument to create 150bp paired-end reads at the FIMM Sequencing unit (Helsinki, Finland). The sequence reads have been deposited to ENA under project PRJEB14064. The accession numbers for individual strains are listed in Table S1. De novo assemblies were performed using velvet ^24^ and annotated using Prokka ^25^. A core genome alignment of the strains was constructed from localised co-linear blocks using the Parsnp tool from the Harvest suite ^26^. A maximum likelihood phylogeny was reconstructed from the alignment using RaxML with 100 bootstrap and the GTR-Gamma model of substitution ^27^. Metadata encompassing information on isolation (continent, country and host), serotype, and CRISPR motif for each strain were superimposed on the tree as colour bars, using the Interactive Tree of Life web-based tool (http://itol.embl.de/) ^28^.

### Analysis of CRISPR loci

The genomic de novo assemblies were searched for CRISPR loci using BLASTN using the *Y. pseudotuberculosis* –specific CRISPR direct repeat sequence (5’- tttctaagctgcctgtgcggcagtgaac-3’), its complementary sequence, the 5’- and 3’-flanking sequences of the YP1, YP2 and YP3 loci and their complementary sequences ^11^. Identified sequences were submitted to the CRISPRFinder tool at CRISPRs Web Server (http://crispr.u-psud.fr/) together with the spacer dictionary compiled earlier ^11^. This analysis increased the number of identified spacers in the *Y. pseudotuberculosis* spacer dictionary from 1902 to 2969 (Table S2). The complete list of the strains and spacer arrays used for CRISPR spacer clustering is in Table S3.

### Accessory genome analysis

The Large Scale Blast Score Ratio (LS-BSR) pipeline ^29^ was used to create pangenomes from genome assemblies of all strains. The included post-matrix script (filter_BSR_variome.py) was run to isolate the accessory genomes from the pangenomes. The resulting accessory genome matrix was then transposed according to the order of the strains on the phylogenetic tree. The output was used to visualise the presence or absence of all accessory genes in each individual genome by generating a heat map using the ggplot2 package of the R statistical software. Genes with >90% prevalence and also those found in fewer than 5 strains were excluded from this analysis. The included Python script compare_BSR.py from LS-BSR was used to look for unique coding sequences (CDSs) between two defined populations in the pangenome matrix. Comparisons were made between the “European” clade of strains and the “Asian” clade, as well as between each CRISPR cluster and the rest of the population. Any unique CDSs detected were compared to the non-redundant nucleotide database using nucleotide BLAST (http://blast.ncbi.nlm.nih.gov/) to determine the genes they encode.

KPAX2 software was used to cluster the strains on the basis of their CRISPR spacer profiles ^30^. Input to the software was a binary matrix with columns representing an absence/presence variable for each of the 2969 spacers in each detected CRISPR cassette. KPAX2 was used with default prior hyperparameters and an upper bound for the number of clusters equal to 50. Five independent runs of the inference algorithm were performed and the clustering solution with the highest posterior probability was chosen. All estimation runs converged to a number of clusters well below the chosen upper bound, indicating that it was sufficiently large to accommodate the region of high posterior density. To analyse the association between CRISPR spacer patterns and the accessory genome content, we calculated an average accessory genome dissimilarity (Hamming distance normalized by the number of CRISPR spacers) matrix for all detected CRISPR clusters with >1 strain (18 clusters).

To assess the significance of the observed dissimilarity pattern, we used a standard permutation test. Under the null hypothesis of no association between CRISPR cluster and the accessory genome content, the cluster label of a strain can be permuted randomly. For each of 10000 random permutations of the labels we then re-calculated the average dissimilarity for each cluster and recorded how often the observed value is smaller than the observed dissimilarity in the original data matrix. Under global significance level of 5% 12/18 CRISPR clusters had a significantly smaller average distance than expected under the null hypothesis.

### Detection of core genome recombination events

Core genome alignments were constructed using Parsnp ^26^. Core genome recombination events were detected by performing BratNextGen analysis on the core genome alignment ^31^. BratNextGen was run using the default prior settings, 20 iterations of the HMM estimation algorithm and 100 runs executed in parallel for the permutation test of significance at 5% level.

### Dating analysis

To date the geographic split within the species, and the formation of the distinct CRISPR clusters, we used BEAST ^32^. The core genome alignment for the 73 strains for which isolation dates were known was obtained using Parsnp, was stripped of recombination detected using BratNextGen, and the resulting alignment used as input with all known dates of isolation to date individual taxa. By assessing ESS scores for priors the following parameters were chosen for the best fitting model: HKY model of substitution with estimated base frequencies and a relaxed molecular clock. The analysis was run for a total of 50 million iterations with the initial 5 million used as burn-in. From this a maximum clade credibility tree was inferred and visualised in Figtree. For the skyline analysis a stepwise constant variant was selected with the age of youngest tip set to zero.

## Acknowledgements

TS is funded by a Nottingham Trent University, Vice Chancellors studentship awarded to TS and AM. JC is partially funded by the COIN Centre of Excellence (Academy of Finland). This work was part of the European Defence Agency (EDA) project B-1325-ESM4-GP. Katarzyna Leskinen, Joanna Zur and Monika Rajtor are thanked for the help in genomic DNA isolation.

---

Figure S1: Maximum likelihood Phylogenetic tree of 134 *Y. pseudotuberculosis* isolates. The phylogeny is derived from a core genome alignment constructed using Parsnp and the phylogenetic tree was visualised using iTOL. The tree is annotated with all relevant meta-data presented in the manuscript.

Figure S2: A) Dated phylogeny of the 73 taxa for which isolation dates exist. The scale bar represents years before present and is annotated with CRISPR clusters as determined by Bayesian clustering of concatenated sequence of CRISPR spacer arrays. The “Asian” and “European” clades are defined by tree branch colouring. B) Bayesian skyline analysis of the population dynamics of the 73 dated taxa. The timeline represents years before present with the median value represented by the solid black line. The shaded area represents 95 confidence interval.

